# Chronic neuronal excitation leads to homeostatic suppression of structural long-term potentiation

**DOI:** 10.1101/2021.03.16.435575

**Authors:** Hiromi H. Ueda, Aiko Sato, Maki Onda, Hideji Murakoshi

## Abstract

Synaptic plasticity is long-lasting changes in synaptic currents and structure. When neurons are exposed to signals that induce aberrant neuronal excitation, they increase the threshold for the induction of synaptic plasticity, called homeostatic plasticity. To further understand the homeostatic regulation of synaptic plasticity and its molecular mechanisms, we investigated glutamate uncaging/photoactivatable (pa)CaMKII-dependent sLTP induction in hippocampal CA1 neurons after chronic neuronal excitation by GABA_A_ receptor antagonists. The neuronal excitation suppressed the glutamate uncaging-evoked Ca^2+^ influx and failed to induce sLTP. Single-spine optogenetic stimulation using paCaMKII also failed to induce sLTP, suggesting that CaMKII downstream signaling is impaired in response to chronic neuronal excitation. Furthermore, while the inhibition of Ca^2+^ influx was protein synthesis-independent, paCaMKII-induced sLTP depended on it. Our findings demonstrate that chronic neuronal excitation suppresses sLTP in two independent ways (i.e., the inhibitions of Ca^2+^ influx and CaMKII downstream signaling), which may contribute to the robust neuronal protection in excitable environments.

## INTRODUCTION

Long-term potentiation (LTP), a form of synaptic plasticity, is a persistent increase in synaptic strength. The molecular mechanism of LTP has been well-studied in the excitatory synapse of the hippocampus (Nicoll, 2017; Yashiro and Philpot, 2008). Presynaptic glutamate binds to postsynaptic N-methyl-D-aspartate (NMDA)-type glutamate receptors (NMDARs). It induces Ca^2+^ influx into the dendritic spines through NMDARs (Yashiro and Philpot, 2008). The increase in Ca^2+^ activates various intracellular signaling molecules such as calmodulin (Bayer and Schulman, 2019; Lisman et al., 2012). The activated calmodulin (Ca^2+^/ calmodulin) binds to Ca^2+^/calmodulin-dependent protein kinase II (CaMKII) (Bayer and Schulman, 2019; Giese and Mizuno, 2013; Herring and Nicoll, 2016; Lisman et al., 2012). It results in the increased kinase activity by the changes of CaMKII structure (Lee et al., 2009; Saneyoshi et al., 2019). The activated CaMKII phosphorylates and recruits signaling molecules (Bosch et al., 2014; Murakoshi and Yasuda, 2012; Nakahata and Yasuda, 2018). These events lead to spine enlargement and α-amino-3-hydroxy-5-methyl-4-isoxazole propionic acid (AMPA)-type glutamate receptors (AMPARs) accumulation in the postsynaptic density (i.e., LTP) (Cingolani and Goda, 2008; Derkach et al., 2007; Malinow and Malenka, 2002). Notably, previous reports have indicated that CaMKII activation is sufficient to trigger LTP (Jourdain et al., 2003; Lledo et al., 1995; Pettit et al., 1994; Shibata et al., 2021). Since previous reports suggest that persistent spine enlargement correlates well with the increase in AMPA currents (i.e., LTP) (Govindarajan et al., 2011; Harvey and Svoboda, 2007; Matsuzaki et al., 2004), this spine enlargement has been termed as structural LTP (sLTP).

The threshold for LTP induction is regulated to maintain optimal neuronal excitability, which is called homeostatic plasticity. More specifically, it is called the sliding threshold, meta-plasticity, or Bienenstock, Cooper, and Munro (BCM) theory (Cooper and Bear, 2012; Keck et al., 2017). In this model, the threshold for LTP induction could increase in response to prolonged neuronal excitation. This suppression of LTP prevents the positive feedback loop of LTP in excitable environments, which contributes to the stabilization of neuronal excitability. For example, after the chronic application of γ-Aminobutyric acid type A (GABA_A_) receptor antagonists in cultured hippocampal slices, high-frequency electrical stimulation of Schaffer collaterals fails to induce LTP (Abegg et al., 2004; Moulin et al., 2019; Suarez et al., 2012). The electrical induction of LTP is also impaired in acute slices after chronic optogenetic excitation administered to the ventral hippocampus of freely moving mice (Moulin et al., 2019). From a physiological point of view, epileptic seizures reduce the LTP accompanying spatial memory deficits (Suarez et al., 2012). These findings demonstrate the functional depression of synaptic plasticity in response to aberrant neuronal excitation. However, whether structural synaptic plasticity is also depressed in excited neurons remains elusive. Furthermore, molecular mechanisms of the homeostatic regulation of synaptic plasticity are poorly understood.

To investigate the occurrence of sLTP, we used two-photon glutamate uncaging (Matsuzaki et al., 2004) and photoactivatable (pa)CaMKII (Shibata et al., 2021) after chronic neuronal excitation in cultured hippocampal slices. We found that chronic neuronal excitation led to the suppression of glutamate uncaging-evoked Ca^2+^ influx into dendritic spines in a protein synthesis-independent manner and failure of sLTP induction. Additionally, the photoactivation of paCaMKII in single spines using two-photon excitation also failed to induce sLTP, but reversed in the presence of protein synthesis inhibitor, suggesting that the chronic excitation impairs CaMKII downstream signaling in a protein synthesis-dependent manner. These results demonstrate that two independent mechanisms (i.e., the inhibitions of Ca^2+^ influx and CaMKII downstream activity) are responsible for robust sLTP suppression after chronic neuronal excitation.

## RESULTS

### Chronic bicuculline application induces neuronal activation and homeostatic depression of spine density

To induce prolonged neuronal excitation, we applied the GABA_A_ receptor antagonist bicuculline (10 μM) to cultured hippocampal slices for 24 hours (Figure 1A). To label the chronically excited neurons, we employed the synthetic activity-dependent promoter, ESARE (Kawashima et al., 2013), in combination with a fast-maturation mutant of yellow fluorescent proteins, Achilles (Yoshioka-Kobayashi et al., 2020) with a destabilization signal (Li et al., 1998), called d2Achilles. We transfected CA1 pyramidal neurons in cultured hippocampal slices by injecting adeno-associated viral vectors (AAVs) encoding ESARE-d2Achilles (Figure 1B). After 5-8 days, we incubated the slices in bicuculline-containing culture media for 24 hours, and successfully observed d2Achilles fluorescence (i.e, chronically excited neurons) (Figure 1C). For the control experiment (no treatment with bicuculline), we injected AAVs-Syn-DIO-Achilles with a low concentration of AAVs-CaMP0.4-Cre to sparsely label the neurons (Figure 1B). Previous studies reported the depression of spine density as a form of homeostatic plasticity in response to neuronal excitation (Fiore et al., 2014; Goold and Nicoll, 2010; Mendez et al., 2018; Moulin et al., 2019). Consistent with this, the spine density of our chronically excited neurons was significantly decreased relative to that of the control neurons (Figures 1D and 1E), indicating that the chronic bicuculline application induces the homeostatic depression of spine density in hippocampal neurons.

**Fig. 1.**
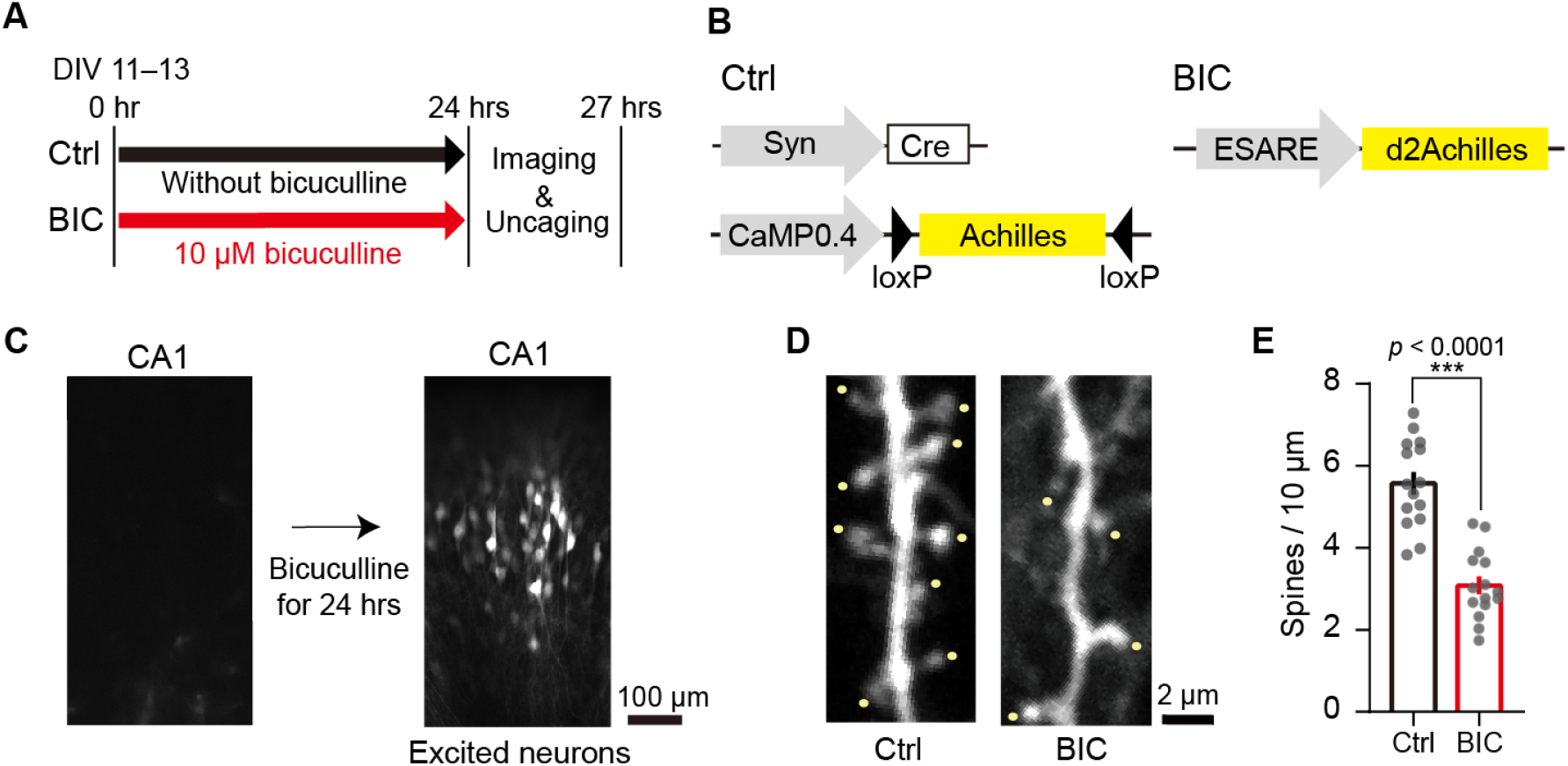
Chronic bicuculline application induces neuronal activation and homeostatic depression of spine density. (A) Schematic timelines of the experimental protocol for two-photon imaging and glutamate uncaging. Hippocampal slices were incubated in a culture medium containing bicuculline (BIC) for 24 hrs to excite the neurons chronically. Subsequently, the slices were placed in an imaging buffer and the experiment was carried out for up to 3 hrs. (B) In the control experiments (Ctrl), AAVs encoding Cre under synapsin promoter (Syn) and Achilles double-floxed inverse orientation (DIO) under a 0.4 kb version of CaMKII promoter (CaMP0.4) were used for sparse labeling. For the group treated with bicuculline (BIC), AAV encoding d2Achilles under the activity-dependent promoter ESARE was used. (C) Epifluorescence images of hippocampal slices transfected with ESARE-d2Achilles. No d2Achilles expression was observed before the treatment with bicuculline. After the treatment, a number of CA1 pyramidal neurons expressed d2Achilles. (D and E) Measurement of spine density in hippocampal CA1 neurons after the treatment with bicuculline. Yellow points indicate counted spines. The number of samples (dendrites/neurons) was 15/5 and 15/5 for Ctrl and BIC, respectively. The data are presented as mean ± standard error of the mean. Statistical comparisons were performed using a two-tailed unpaired *t* test. \*\*\**p* < 0.001.

### Glutamate uncaging fails to induce sLTP in chronically excited neurons

To investigate the structural plasticity of dendritic spines in chronically excited neurons, we applied a low-frequency train of two-photon glutamate uncaging at a spine and monitored Achilles fluorescence of the spine by two-photon excitation at 920 nm (Figures 2A and 2B). In the control experiments, the volume of the stimulated spines, but not adjacent spines, rapidly increased (317%, 4-6 min: transient phase) and relaxed to an elevated volume for 20-30 min (133%, 20-30 min: sustained phase) (Figures 2C and 2D). Contrastingly, the enlargement of the stimulated spines did not occur in chronically excited neurons (Figures 2A-2D). After chronic neuronal excitation by another GABA_A_ receptor antagonist, gabazine, glutamate uncaging also failed to induce sLTP (Figures 2E-2H). These results demonstrate that chronic neuronal excitation leads to the depression of sLTP in hippocampal CA1 neurons.

**Fig. 2.**
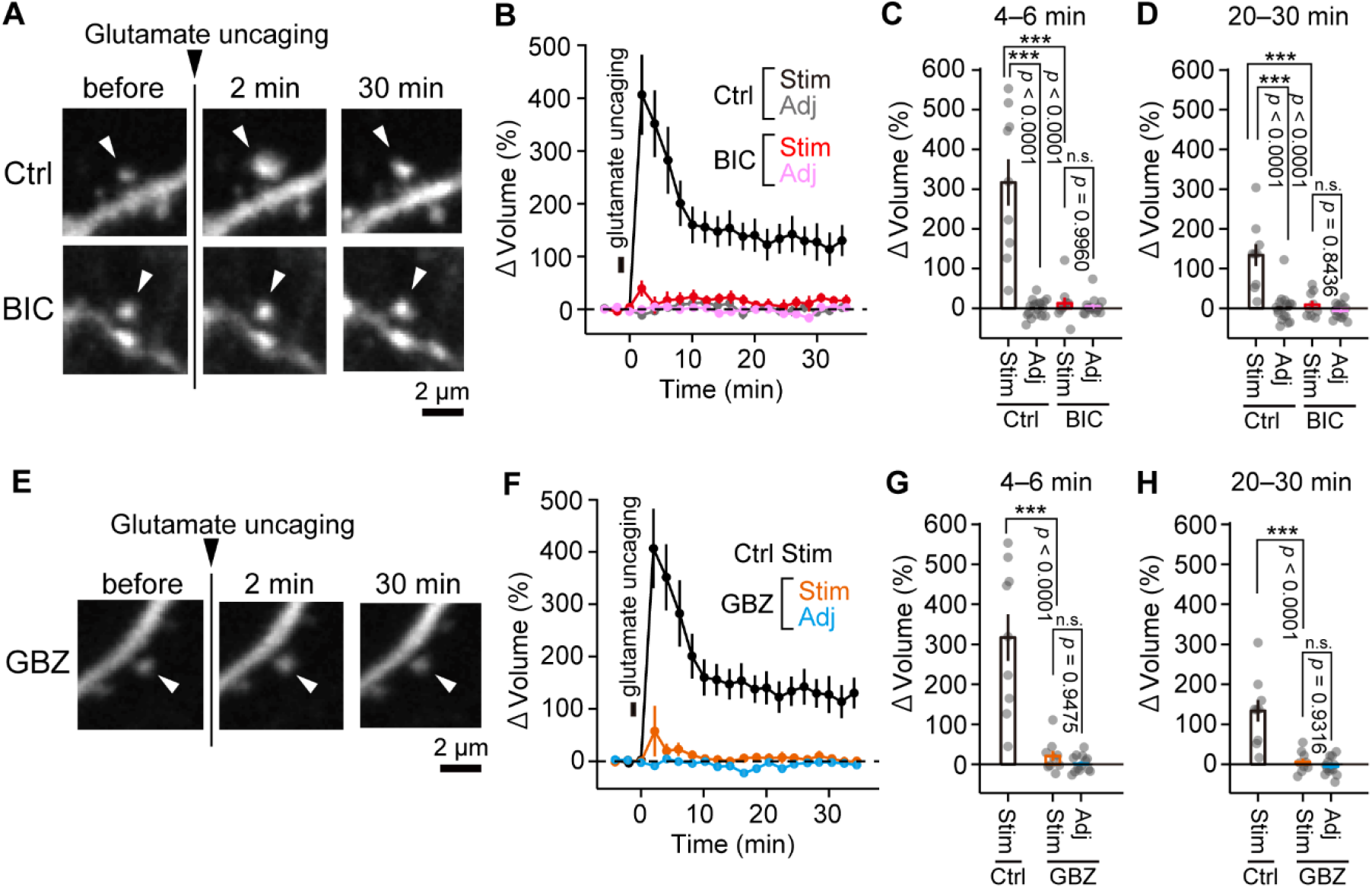
Glutamate uncaging fails to induce sLTP in chronically excited neurons. (A and E) Two-photon fluorescence images of dendritic spines during the induction of sLTP by two-photon glutamate uncaging. Hippocampal CA1 neurons expressing Achilles or d2Achilles were observed by two-photon excitation at 920 nm, and MNI-glutamate was uncaged at 720 nm (30 trains, 0.5 Hz, 6 ms duration/pulse, 6 mW) on a spine indicated by white arrows. (B) Averaged time course of the change in spine volume upon glutamate uncaging after treatment with bicuculline in the stimulated (BIC-Stim) and adjacent spines (2-10 μm, BIC-Adj). For comparison, the time course of the stimulated spines (Ctrl-Stim) and adjacent spines (2-10 μm, Ctrl-Adj) that were not treated with bicuculline is also shown. The number of samples (spines/neurons) was 9/6 for Ctrl-Stim, 18/9 for Ctrl-Adj, 9/5 for BIC-Stim, and 14/5 for BIC-Adj. (C and D) Quantification of the transient (C, averaged over 4-6 min) and sustained (D, averaged over 20-30 min) change in spine volume. The data are presented as mean ± standard error of the mean (SEM). Statistical comparisons were performed using one-way analysis of variance followed by Turkey’s test. ****p < 0.0001; n.s. represents p > 0.05. (F) Averaged time course of the change in spine volume upon glutamate uncaging after gabazine treatment in the stimulated (GBZ-Stim) and adjacent spines (2-10 μm, GBZ-Adj). For comparison, the time course of the stimulated spines (Ctrl-Stim in Fig. 2B) with no treatment is replotted. The number of samples (spines/neurons) was 10/4 for GBZ-Stim, and 13/4 for GBZ-Adj. (G and H) Quantification of the transient (G, averaged over 4-6 min) and sustained (H, averaged over 20-30 min) change in spine volume. The data are presented as mean ± SEM. Statistical comparisons were performed using one-way analysis of variance followed by Tukey’s test. ***p < 0.001; n.s. represents p > 0.05.

### The inhibition of protein synthesis partially recovers sLTP

Prolonged neuronal excitation induces protein synthesis and degradation (Dorrbaum et al., 2020; Schanzenbacher et al., 2018; Schanzenbacher et al., 2016). It has been reported that protein synthesis is required for homeostatic depression of spine density (Mendez et al., 2018). Thus, we expected that sLTP suppression would depend on protein synthesis. To test this, we applied the protein synthesis inhibitor anisomycin (100 μM) along with bicuculline to hippocampal slices for 24 hours (Figure 3A). Since anisomycin inhibits activity-dependent d2Achilles protein synthesis, we injected the AAVs-CaMP0.4-DIO-Achilles with a low concentration of AAVs-Syn-Cre, instead of AAVs-ESARE-d2Achilles (Figure 3B). We confirmed that glutamate uncaging-induced sLTP was also impaired in Achilles-expressing neurons after the bicuculline treatment (Figures 3C-3F). Furthermore, the application of anisomycin partially recovered sLTP (Figures 3C-3F). These results demonstrate that the suppression of sLTP is partially dependent on protein synthesis.

**Fig. 3.**
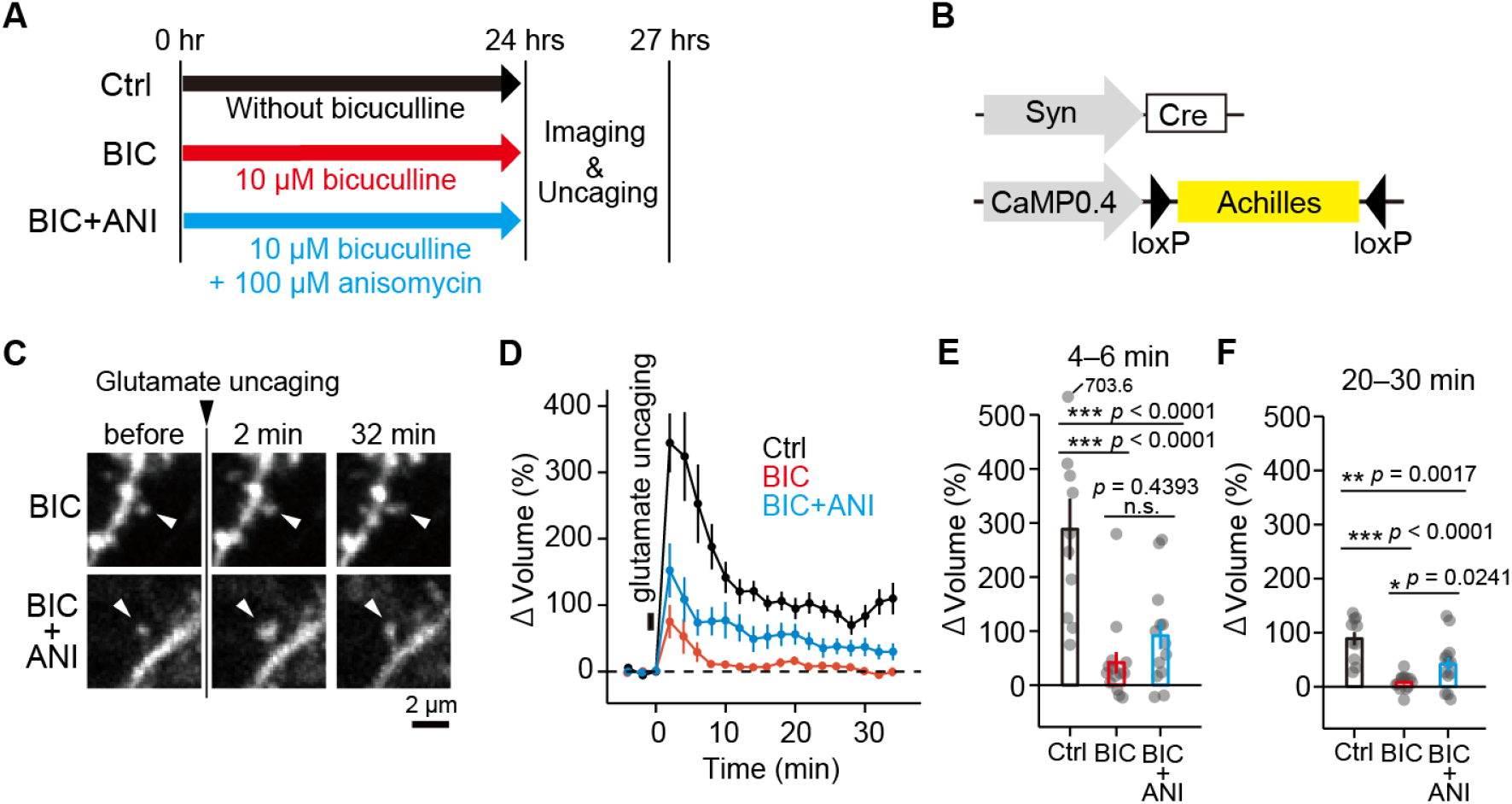
The inhibition of protein synthesis partially recovers sLTP. (A) Schematic timelines of control (Ctrl), bicuculline treatment (BIC), and anisomycin with bicuculline treatment (BIC+ANI). (B) Schematic of AAV-Syn-Cre and AAV-CaMP0.4-DIO-Achilles used in the experiments. (C) Two-photon fluorescence images of dendritic spines during the induction of sLTP by two-photon glutamate uncaging. A hippocampal CA1 neuron expressing Achilles was observed by two-photon excitation at 920 nm, and caged glutamate was uncaged at 720 nm (30 trains, 0.5 Hz, 6 ms duration/pulse, 6 mW) on a spine indicated by white arrows. (D) Averaged time course of the change in spine volume in stimulated spines in the control condition, bicuculline treatment, and anisomycin with bicuculline treatment. The number of samples (spines/neurons) was 10/7 for Ctrl, 14/7 for BIC, and 13/5 for BIC+ANI. (E and F) Quantification of the transient (E, averaged over 4-6 min) and sustained (F, averaged over 20-30 min) change in spine volume. The data are presented as mean ± standard error of the mean. Statistical comparisons were performed using one-way analysis of variance followed by Tukey’s test. \*\*\**p* < 0.001; \*\**p* < 0.01; \**p* < 0.05; n.s. represents *p* > 0.05.

### Glutamate uncaging-induced Ca^2+^ influx into single spines decreases after chronic neuronal excitation

Chronic neuronal excitation has been shown to induce the depression of NMDAR currents (Goold and Nicoll, 2010; Watt et al., 2000). Therefore, it is possible that the chronic excitation may depress NMDAR-dependent Ca^2+^ influx, resulting in the suppression of sLTP. To measure the Ca^2+^ influx, we transfected CMV-GCaMP6f-P2A-mScarlet into CA1 pyramidal neurons using a gene gun and monitored the GCaMP6f transient in a spine after a single pulse of glutamate uncaging (720 nm, 6 ms duration/pulse, 6 mW) in the presence and absence of bicuculline and anisomycin (Figures 4A-4C) (Chen et al., 2013).

**Fig. 4.**
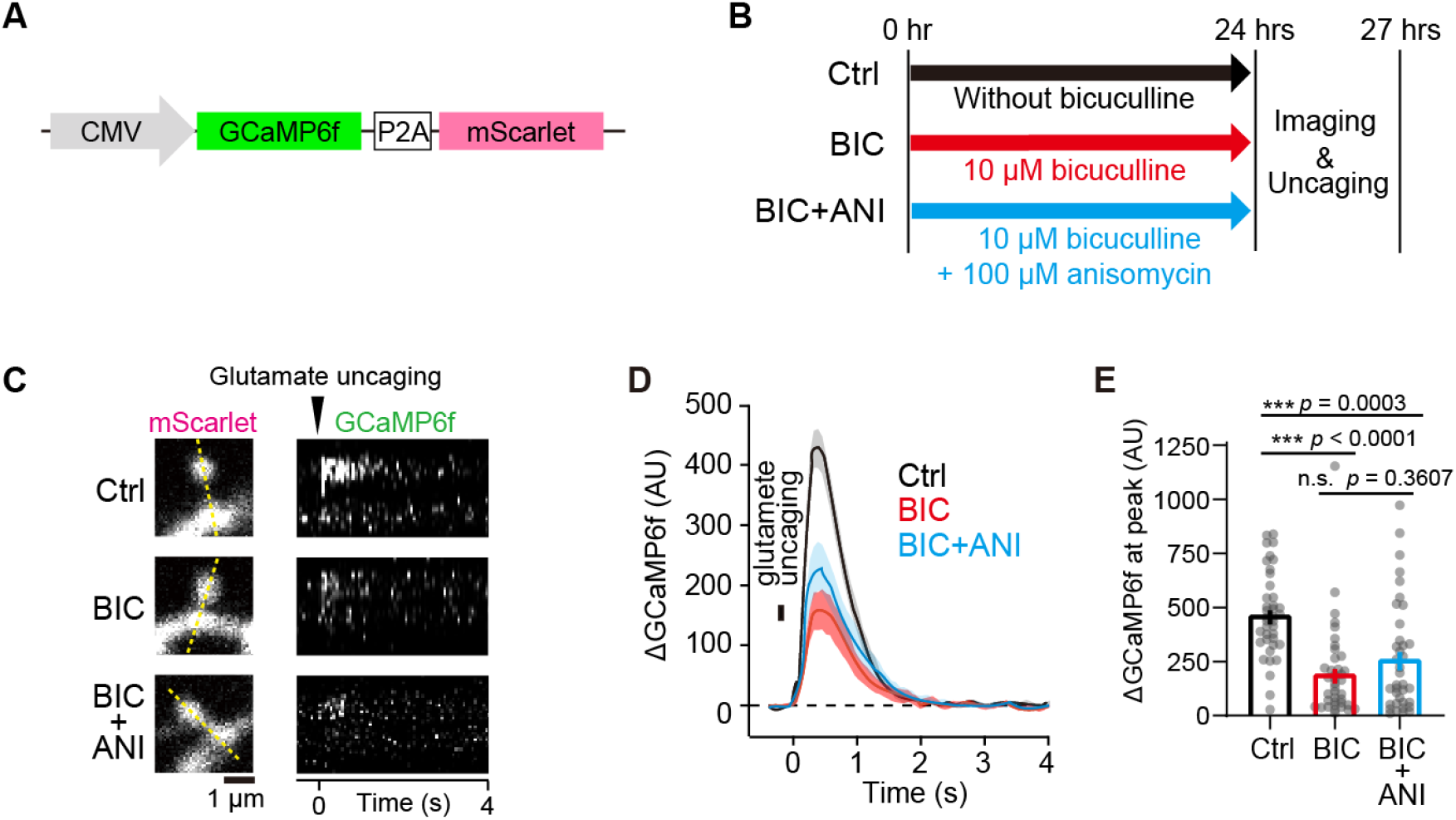
Glutamate uncaging-induced Ca^2+^ influx into single spines decreases after chronic neuronal excitation. (A) Schematic of CMV-GCaMP6f-P2A-mScarlet transfected into hippocampal CA1 neurons. (B) Schematic timelines of control (Ctrl), bicuculline treatment (BIC), and anisomycin with bicuculline (BIC+ANI) treatment. (C) Two-photon fluorescence images of the spines (mScarlet) before glutamate uncaging (left column) and GCaMP6f fluorescence of Ca^2+^ transients in the spines after uncaging (right column), which is shown as a kymograph of yellow dash lines. Both mScarlet and GCaMP6f were imaged by two-photon excitation at 1000 nm, and caged glutamate was uncaged at 720 nm (1 trains, 6 ms duration/pulse, 6 mW) at the tip of the spines. (D) Averaged time course of Ca^2+^ change in the stimulated spines in control; 24 hrs bicuculline treatment; bicuculline and anisomycin treatment. The number of samples (spines/neurons) was 39/10 for Ctrl, 39/11 for BIC, and 37/9 for BIC+ANI. (E) Quantification of peak changes of Ca^2+^ transients. The data are presented as mean ± standard error of the mean. Statistical comparisons were performed using one-way analysis of variance followed by Tukey’s test. \*\*\**p* < 0.001; n.s. represents*p* > 0.05.

We found that uncaging-evoked Ca^2+^ transients decreased in the neurons treated with bicuculline (Figure 4C). Quantitative analysis revealed that the peak amplitude of the Ca^2+^ transients was significantly lower in the bicuculline treatment than that of the Ca^2+^ transients in the control neurons (Figures 4D and 4E). The application of anisomycin with bicuculline did not reverse the suppression of the Ca^2+^ transients (Figures 4C-4E), suggesting that the suppression of the Ca^2+^ influx is not dependent on protein synthesis. This result is consistent with the findings from a previous study wherein NMDAR currents were depressed via a protein synthesis-independent pathway after chronic neuronal excitation (Goold and Nicoll, 2010). Conversely, sLTP suppression was partially dependent on protein synthesis (Figure 3). Thus, our findings suggest that other mechanisms may be involved in sLTP depression besides the suppression of Ca^2+^ transients.

### paCaMKII activation fails to induce sLTP in chronically excited neurons

Next, we investigated the downstream of Ca^2+^ in the signal cascade for sLTP. It has been shown that the activation of CaMKII downstream signaling is necessary to induce sLTP in hippocampal neurons (Bayer and Schulman, 2019; Giese and Mizuno, 2013; Herring and Nicoll, 2016; Lisman et al., 2012). Thus, it is possible that the impairment of CaMKII downstream signaling causes the suppression of sLTP. To examine this hypothesis, we directly activated CaMKII downstream signaling by using the genetically encoded paCaMKII (Figure 5A) (Shibata et al., 2021). Two-photon excitation of paCaMKII enables the activation of CaMKII downstream molecules in single spines and induces sLTP without Ca^2+^ influx (Shibata et al., 2021). We co-transfected hippocampal neurons by injecting AAVs encoding tdTomato-P2A-paCaMKII with ESARE-d2Achilles or AAVs in control experiments (Figure 5B). First, we identified the neurons expressing paCaMKII by observing tdTomato fluorescence using epifluorescence microscopy. Subsequently, we monitored the neurons by observing d2Achilles/Achilles fluorescence using two-photon microscopy at an excitation wavelength of 1010 nm (Figure 5C). To induce paCaMKII activation in single spines, we applied a low-frequency train of two-photon excitation pulses to single spines (820 nm, 30 pulses, 0.5 Hz, 80 ms duration/pulse, 4 mW) (Figure 5B). In the control experiments, the spine volume increased rapidly by approximately 301% following paCaMKII activation (4-6 min) and relaxed to an elevated level of 91% during 20-30 min (Figures 5D-5F). By contrast, chronically excited neurons did not show spine enlargement (Figures 5D-5F), similar to the results of glutamate uncaging (Figures 2A-2D). These results indicate that paCaMKII-induced sLTP was suppressed in chronically excited neurons. A possible explanation for this suppression is the difference in the expression/activity of paCaMKII between the chronically excited and control neurons; to investigate this, we examined the expression level and activity of paCaMKII using a biochemical assay. We transfected the dissociated hippocampal neurons with CaMK0.4-paCaMKII and ESARE-mScarlet using AAVs (Figure 5G), and confirmed neuronal activation after the treatment with bicuculline (10 μM) by monitoring the robust expression of mScarlet (Figure 5H). We evaluated the expression and pT286 autophosphorylation of paCaMKII under blue light illumination. Chronic neuronal excitation did not change the expression or light-induced activation of paCaMKII (Figure 5H). Thus, paCaMKII-induced sLTP may be suppressed due to the inhibition of CaMKII downstream rather than paCaMKII itself.

**Fig. 5.**
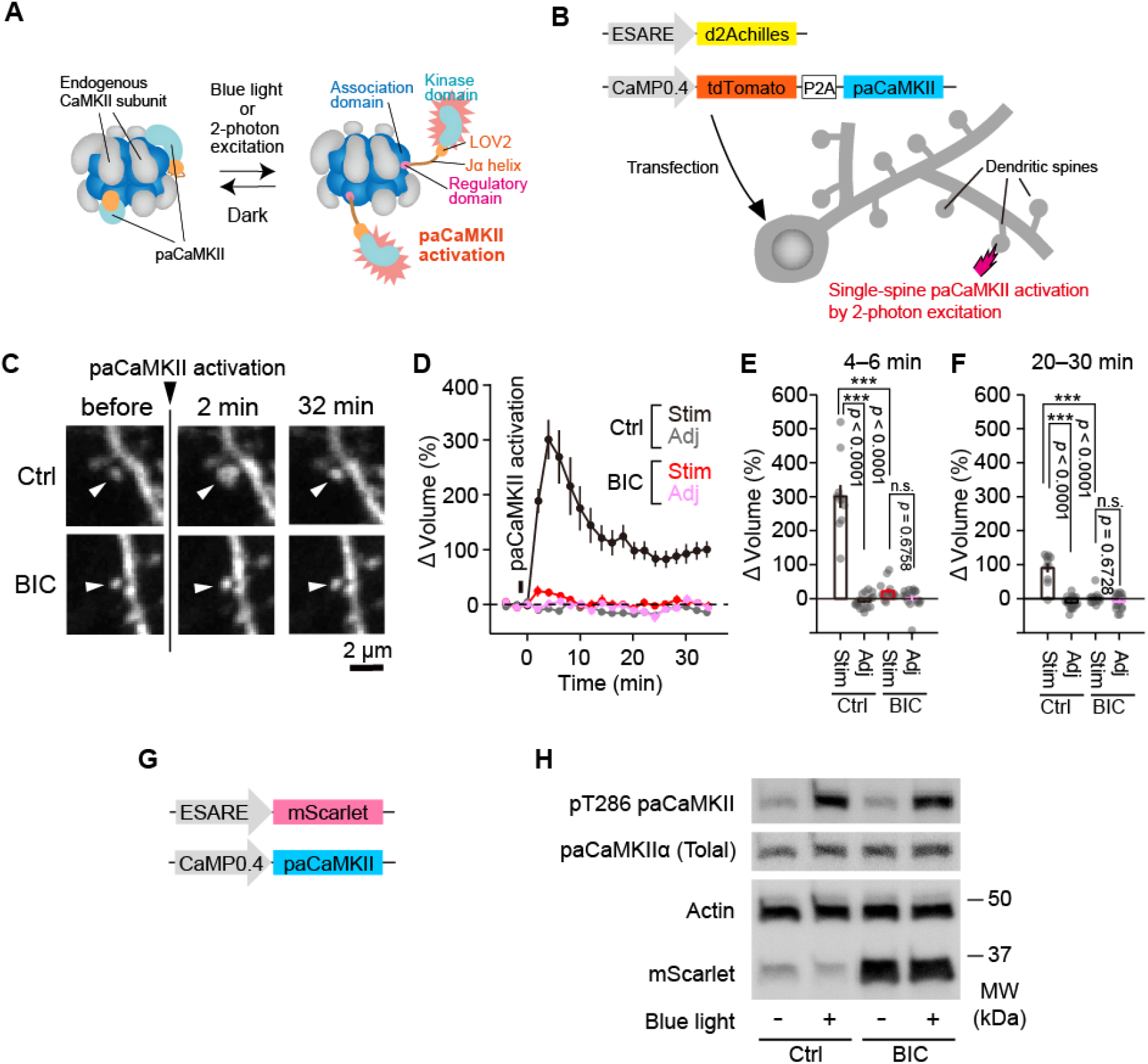
paCaMKII activation fails to induce sLTP in chronically excited neurons. (A) Schematic drawing of paCaMKII activation in the oligomeric state. Two-photon excitation induces a structural change of paCaMKII, thereby activating it. Note that paCaMKII can be integrated into an endogenous CaMKII oligomer. The figure was adopted from a previous study (Shibata et al., 2021). (B) Schematic representation of sLTP induction by paCaMKII activation. AAVs encoding CaMP0.4-tdTomato-P2A-paCaMKII and ESARE-d2Achilles were co-transfected into the neurons. (C) Two-photon fluorescence images of dendritic spines during the induction of sLTP by two-photon paCaMKII activation. Hippocampal CA1 neurons expressing Achilles or d2Achilles and tdTomato-P2A-paCaMKII were observed by two-photon excitation at 1010 nm, and paCaMKII was activated at 820 nm (30 trains, 0.5 Hz, 80 ms duration/pulse, 4 mW) in the spine indicated by white arrows. (D) Averaged time course of the change in spine volume in the stimulated spine (BIC-Stim) and adjacent spines (2–10 μm, BIC-Adj) 24 hrs after bicuculline application. A control experiment (Ctrl-Stim) and adjacent spines (2–10 μm, Ctrl-Adj) are also shown. The number of samples (spines/neurons) was 10/4 for Ctrl-Stim, 19/4 for Ctrl-Adj, 14/5 for BIC-Stim, and 17/5 for BIC-Adj, respectively. (E and F) Quantification of the transient (C, averaged over 4–6 min) and sustained (D, averaged over 20–30 min) change in spine volume. Data are presented as the mean ± standard error of the mean. Statistical comparisons were performed using one-way analysis of variance followed by Tukey’s test. \*\*\**p* < 0.001; n.s. represents *p* > 0.05. (G) Schematic of AAV encoding ESARE-mScarlet and CaMP0.4-paCaMKII transfected into neurons for biochemical assays. (H) Dissociated hippocampal neurons expressing paCaMKII were illuminated with blue light for 5 min after the treatment with bicuculline (lane 1: no bicuculline/no light; lane 2: no bicuculline/with light; lane 3: bicuculline/no light; lane 4: bicuculline/with light). The kinase activity of paCaMKII and the expression of paCaMKII, actin proteins, and mScarlet were evaluated by western blotting.

### The activity of the CaMKII pathway is not saturated in chronically excited neurons

The application of bicuculline induces an increase in the intracellular Ca^2+^ concentration in cultured hippocampal slices (van der Linden et al., 1993). Since Ca^2+^ activates CaMKII pathways in dendritic spines, the long-lasting Ca^2+^ increase induced by the chronic application of bicuculline may saturate the activity of CaMKII downstream. To examine whether the CaMKII downstream activity is saturated, we augmented paCaMKII activation by increasing the duration per pulse of the two-photon excitation with a fixed pulse number. In the neurons with no bicuculline treatment, extended activation (320 ms/pulse) induced a large spine enlargement compared to that observed in the control stimulation (80 ms/pulse) (Figure 6). This is most likely due to the increase in the activity of CaMKII downstream signaling. In chronically excited neurons, prolonged activation of paCaMKII (320 ms/pulse) successfully induced sLTP (Figure 6), indicating that CaMKII downstream molecules were activated in chronically excited neurons. These results suggest that chronic neuronal excitation makes the activity of CaMKII downstream suppressed rather than saturated.

**Fig. 6.**
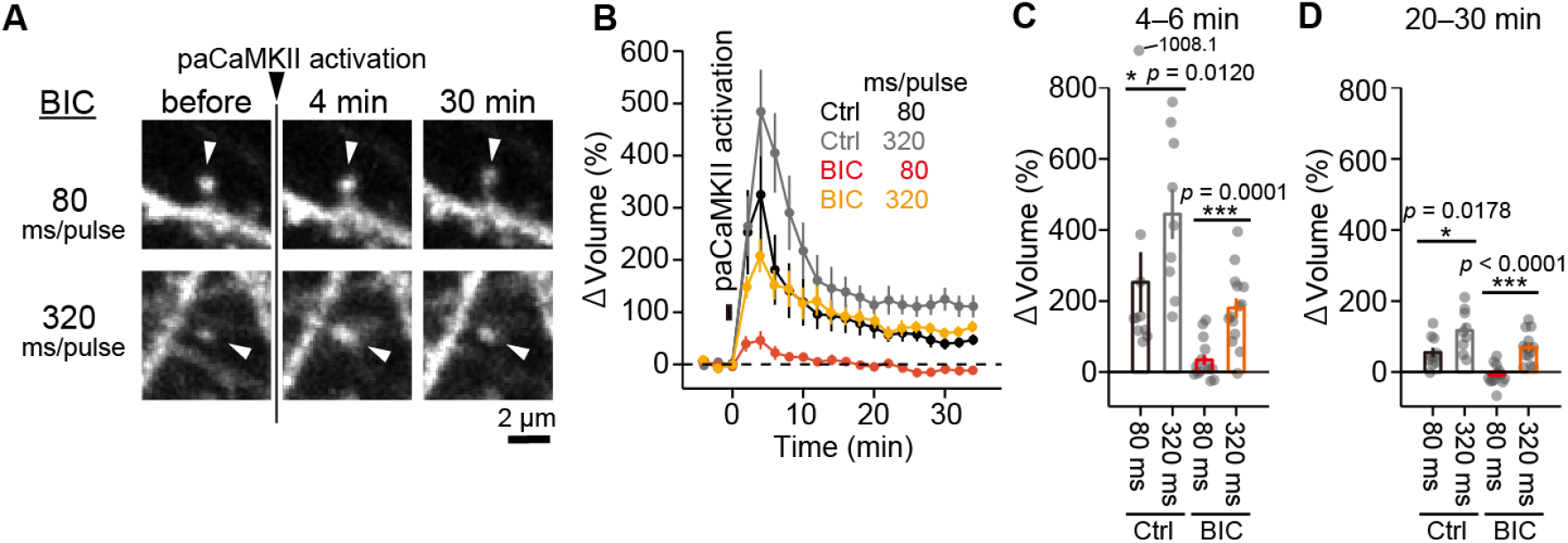
The activity of CaMKII pathway is not saturated in chronically excited neurons. (A) Two-photon fluorescence images of dendritic spines during the induction of sLTP by two-photon paCaMKII activation after the bicuculline treatment. A hippocampal CA1 neuron expressing Achilles or d2Achilles and tdTomato-P2A-paCaMKII was observed by two-photon excitation at 1010 nm, and paCaMKII was activated at 820 nm (30 trains, 0.5 Hz, 80 ms or 320 ms duration/pulse, 4 mW) in a spine indicated by white arrows. (B) Averaged time course of the change in spine volume after paCaMKII activation with different pulse durations (80 or 320 ms/pulse) are plotted for both the control (Ctrl) and bicuculline-treated neurons (BIC). The number of samples (spines/neurons) was 10/5 for Ctrl with 80 ms/pulse, 9/3 for Ctrl with 320 ms/pulse, 14/5 for BIC with 80 ms/pulse, and 14/5 for BIC with 320 ms/pulse. (C and D) Quantification of the transient (C, averaged over 4–6 min) and sustained (D, averaged over 20–30 min) change in spine volume. The data are presented as mean ± standard error of the mean. Statistical comparisons were performed using one-way analysis of variance followed by Tukey’s test. \*\*\**p* < 0.001; \**p* < 0.05;.

### The suppression of paCaMKII-induced sLTP is dependent on protein synthesis

Finally, we examined whether the suppression of paCaMKII-induced sLTP requires the newly synthesized proteins during chronic neuronal activation. We transfected AAVs-tdTomato-P2A-paCaMKII and AAVs-CaMP0.4-DIO-Achilles with a low amount of AAVs-CaMP0.4-Cre for sparse labeling. We confirmed that paCaMKII-induced sLTP was also impaired in Achilles-expressing neurons after the bicuculline treatment (Figure 7). Notably, the inhibition of sLTP was reversed by the inhibition of protein synthesis (Figure 7). These results suggest that sLTP inhibition may be caused by the downstream inhibition of CaMKII by newly synthesized proteins.

**Fig. 7.**
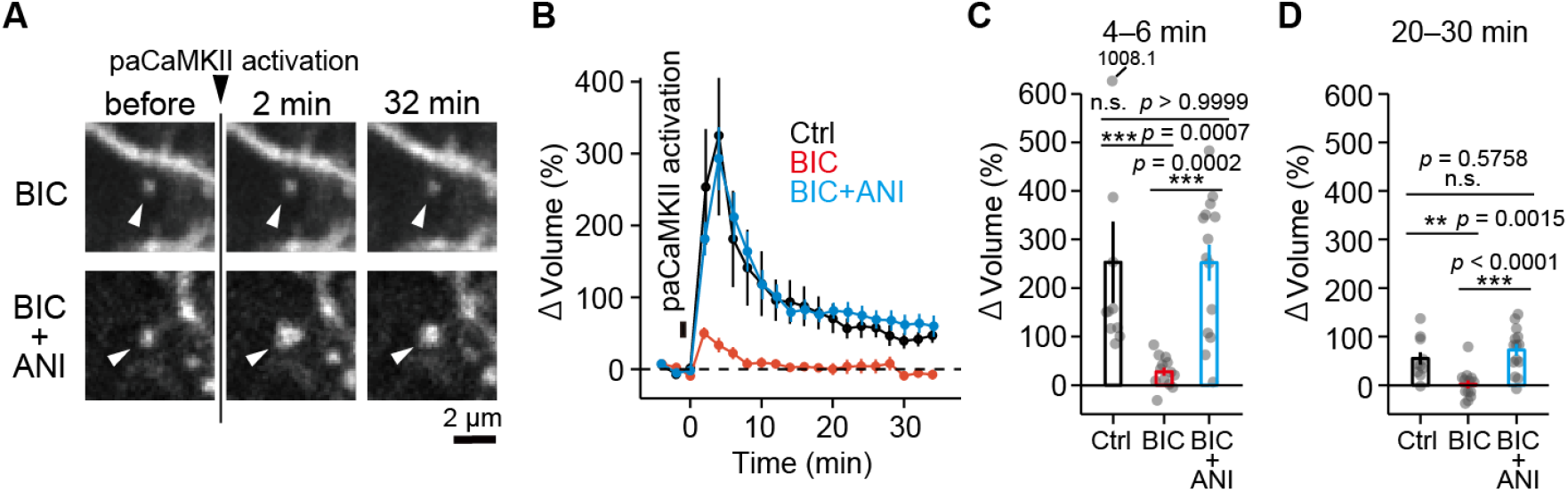
The suppression of paCaMKII-induced sLTP is dependent on protein synthesis. (A) Two-photon fluorescence images of dendritic spines during the induction of sLTP by two-photon paCaMKII activation. A hippocampal CA1 neuron expressing Achilles and tdTomato-P2A-paCaMKII was observed by two-photon excitation at 1010 nm, and paCaMKII was activated at 820 nm (30 trains, 0.5 Hz, 80 ms duration/pulse, 4 mW) in a spine indicated by white arrows. (B) Averaged time course of the change in spine volume after paCaMKII activation after the bicuculline (BIC) or anisomycin (ANI) with bicuculline (BIC+ANI) treatment. For comparison, the averaged timecourse of control experiments (no bicuculline/anisomycin treatment) is also replotted (Ctrl). The number of samples (spines/neurons) was 10/5 for Ctrl, 13/4 for BIC, and 14/5 for BIC+ANI. (C and D) Quantification of the transient (C, averaged over 4–6 min) and sustained (D, averaged over 20–30 min) change in spine volume. The data are presented as mean ± standard error of the mean. Statistical comparisons were performed using one-way analysis of variance followed by Tukey’s test. \*\*\**p* < 0.001; \*\**p* < 0.01; n.s. represents *p* > 0.05.

## DISCUSSION

In this study, we demonstrated that sLTP induction is suppressed by chronic neuronal excitation in hippocampal neurons, most likely via the inhibition of Ca^2+^ influx into dendritic spines and the inhibition of CaMKII downstream pathways. While the downstream inhibition of CaMKII is dependent on protein synthesis, the inhibition of Ca^2+^ influx is protein synthesis-independent. Thus, this two-step inhibitory mechanism may contribute to the robust inhibition of sLTP to stabilize the synaptic structure and excitability.

We found that glutamate uncaging-induced Ca^2+^ influx was suppressed after chronic neuronal excitation. It has been shown that the Ca^2+^ permeability of dendritic spines depends on the subunit composition of postsynaptic NMDARs (Lee et al., 2010; Sobczyk et al., 2005). Glutamate uncaging-induced Ca^2+^ influx through NR2B subunit-containing NMDARs is higher than that through NR2A subunit-containing NMDARs (Sobczyk et al., 2005). Some studies have shown that the expression of NR2B-containing NMDARs decreases after chronic neuronal excitation (Ehlers, 2003; Perez-Otano and Ehlers, 2005; Schanzenbacher et al., 2016). Thus, the decrease in Ca^2+^ influx may be caused by the downregulation of NR2B-containing NMDARs in dendritic spines. Accompanied by the suppression of Ca^2+^ influx, glutamate uncaging-induced sLTP was suppressed after the chronic neuronal excitation. Previous studies have proposed that the threshold of LTP is adjusted in response to neuronal excitation, a phenomenon termed as the sliding threshold model (Cooper and Bear, 2012; Keck et al., 2017). It is well established that the composition of postsynaptic NMDARs is a key determinant of the threshold for LTP, because the Ca^2+^ influx through the receptors triggers LTP. (Cooper and Bear, 2012; Keck et al., 2017; Lee et al., 2010). The suppression of Ca^2+^ influx and sLTP in our results supports the NMDAR-dependent mechanism of the sliding threshold model.

While the change in NMDAR composition is a well-established molecular mechanism of the sliding threshold model, other additional mechanisms remain unexplored. Here, we induced sLTP by the direct activation of CaMKII signaling using paCaMKII. It has been shown that the activation of paCaMKII induces LTP without Ca^2+^ influx (Shibata et al., 2021). We found that paCaMKII-induced sLTP was suppressed by chronic neuronal excitation. Since the expression and function of paCaMKII did not change after the chronic excitation, the downstream activity of CaMKII could have been impaired. The augmentation of paCaMKII activation successfully induced sLTP even in chronically excited neurons, implying that the chronic excitation increases the threshold for activation of CaMKII downstream signaling. These results demonstrate that there must be an NMDAR-Ca^2+^-independent mechanism that explains the increased threshold for sLTP.

We also found that the impairment of sLTP, especially paCaMKII-induced sLTP, is protein synthesis-dependent. This suggests that newly synthesized proteins during the neuronal excitation inhibit the CaMKII downstream signaling for sLTP. In fact, it has been shown that numerous proteins are expressed after the chronic application of bicuculline (Schanzenbacher et al., 2018; Schanzenbacher et al., 2016), and they are known to regulate the shape and size of dendritic spines. For example, since Homer1a, Plk2, and Nr4a1 are expressed by the chronic application of GABA_A_ receptor antagonists and induce homeostatic depression of the spine density, these proteins could be candidate proteins for sLTP inhibition (Chen et al., 2014; Fiore et al., 2014; Hu et al., 2010; Lee et al., 2011; Sala et al., 2003; Seeburg and Sheng, 2008).

Based on our findings, we propose that the homeostatic depression of the CaMKII pathway coupled with a decrease in Ca^2+^ influx is a mechanism for the sliding threshold model (Figure 8). To the best of our knowledge, this is the first report showing the NMDAR-independent mechanism of the sliding threshold model. These two mechanisms for sLTP suppression possibly complement each other to maintain reliable homeostatic plasticity and the stabilization of neuronal excitability.

**Fig. 8.**
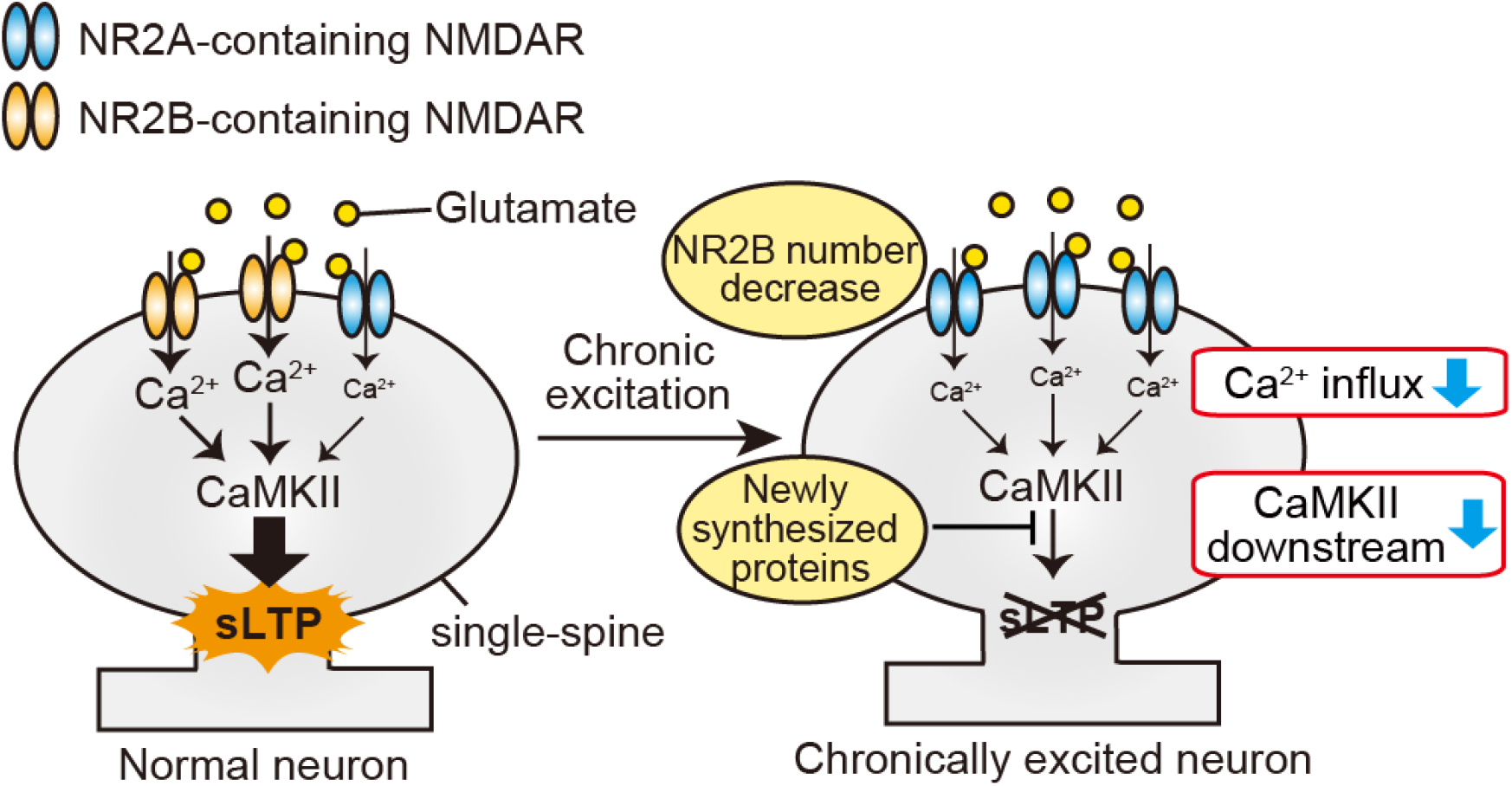
Schematic model of homeostatic suppression of sLTP. Glutamate binds to NMDARs, leading to Ca^2+^ influx into a single spine. Ca^2+^ activates CaMKII signaling, resulting in sLTP induction. Contrastingly, Ca^2+^ influx and CaMKII downstream signaling are depressed in chronically activated neurons. The suppression of Ca^2+^ influx is due to the protein synthesis-independent downregulation of NMDARs. The inhibition of CaMKII downstream signaling is protein synthesis-dependent. These two independent inhibitory mechanisms may contribute to the robust homeostatic suppression of sLTP.

## ACKNOWLEDGEMENTS

This work was supported in part by a Grant-in-Aid for Scientific Research in Innovative Areas (18H02708, 18K19382, 18H04748 Resonance Bio, 19H05434 Singularity Biology, and 16K15225 and JP16H06280 Advanced Bioimaging Support (ABiS) to H.M.) from the MEXT/Japan Society for the Promotion of Sciences (JSPS), Core Research for Evolutional Science and Technology (CREST) (to H.M.) from the Japan Science and Technology Agency (JST), the Research Foundation for Opto-Science and Technology (to H.M.), the Japan Foundation for Applied Enzymology (to H.M.), the Takeda Science Foundation, the Asahi Glass Foundation (to H.M.), Frontier Photonic Sciences Project of National Institutes of Natural Sciences (NINS).

## AUTHOR CONTRIBUTIONS

H.H.U. and H.M. conceived and designed the project. H.H.U. performed the experiments and analyzed the data. A.S., M.O., and H.M. contributed to the cell culture preparation, reagent preparation, and software analysis. H.H.U. and H.M. wrote the manuscript. All the coauthors discussed the results and exchanged comments on the manuscript.

## DECLARATION OF INTERESTS

The authors declare no competing interests.

## METHODS

### Animals

All the animal procedures were approved by the National Institute of Natural Sciences Animal Care and Use Committee, and were performed in accordance with the relevant guidelines and regulations. All the slice cultures were prepared using C57BL/6N mice (SLC, Shizuoka, Japan). This study used dissociated and slice cultures from both male and female pups.

### Reagents

Bicuculline was purchased from Wako Pure Chemical Industries (Osaka, Japan). SR95531 (gabazine) and 4-methoxy-7-nitroindolinyl-caged-L-glutamate (MNI-caged glutamate) were purchased from Tocris Bioscience (Bristol, UK). Anisomycin was purchased from Sigma-Aldrich (St. Louis, MO, USA).

### Plasmids

Plasmids containing *CaMKIIα, ESARE/d2Venus*, and *CaMKII0.4* promoter genes were gifts from Y. Hayashi, H. Bito, and M. Ehlers, respectively. Plasmids containing *WPRE3, Cre, GCaMP6f*, and hSyn-DIO-EGFP genes were gifts from Bong-Kiun Kaang, Connie Cepko, D. Kim, and Bryan Roth (Addgene plasmid #61463, #13775, #40755, #50457), respectively. The synthesized gene encoding *the d2Achilles* gene was purchased from FASMAC (Atsugi, Japan). pAAV-RC-DJ (AAV2/DJ) and pAAV-MCS/pAAV-Helper were purchased from Cell Biolabs (San Diego, CA, USA) and Agilent Technologies (Santa Clara, CA, USA), respectively.

The plasmids, namely ESARE-d2Achilles-SV40polyA, ESARE-d2Venus-SV40polyA, Syn-Cre-WPRE, CaMP04-tdTomato-P2A-paCaMKIIα-WPRE3, and CaMP0.4-DIO-Achilles-WPRE3 were constructed by inserting the respective components into the pAAV-MCS. The CMV-GcaMP6f-P2A-mScarlet plasmid was constructed by inserting GcaMP6f (Chen et al., 2013) and mScarlet (Bindels et al., 2017), together with the P2A (Donnelly et al., 2001) sequence ATNFSLLKQAGDVEENPGP into the modified pEGFP-C1 vector by replacing EGFP (Clontech).

### AAV production and purification

The preparation of AAVs has been described previously in detail (Lock et al., 2010; Shibata et al., 2021). Briefly, HEK293 cells were transfected with the plasmids in a 1:1.6:1 ratio (45 μg of a transgene in pAAV, 72 μg of pAAV-Helper, and 45 μg of pAAV-RC-DJ [AAV2/DJ]) using the polyethylenimine method (Lock et al., 2010). Subsequently, the dishes were incubated at 37 °C and 5% CO_2_ for 96 hrs. The collected culture medium was centrifuged and filtered to remove the cell debris. The clarified supernatant containing the AAV was concentrated using a cross-flow cassette (Vivaflow 50, 100,000MWCO, Sartorius; Goettingen, Germany) or Amicon Ultra-15 (100,000MWCO, Merck, Kenilworth, NJ, USA). Iodixanol step gradients were performed as described by Addgene (homepage section: AAV purification by iodixanol gradient ultracentrifugation). The buffer solution of the virus was exchanged with phosphate-buffered saline at different concentrations. The titer of AAVs was determined by quantitative polymerase chain reaction using the THUNDERBIRD qPCR Mix (Toyobo, Osaka, Japan). The resultant virus titers typically ranged between 2 × 10^9^ and 2 × 10^10^ genome copies/μL in a total volume of ~400 μL.

### Organotypic hippocampal slices and gene transfection by AAV or gene gun

Hippocampal slices were prepared from postnatal day 6–9 C57BL/6N mice as described previously (Stoppini et al., 1991). Briefly, the animal was deeply anesthetized with isoflurane, after which the animal was quickly decapitated and the brain removed. The hippocampi were isolated and cut into 350 μm sections in an ice-cold dissection medium (250 mM N-2-hydroxyethylpiperazine-N’-2-ethanesulfonic acid, 2 mM NaHCO_3_, 4 mM KCl, 5 mM MgCI_2_, 1 mM CaCl_2_, 10 mM D-glucose, and 248 mM sucrose). The slices were cultured on the membrane inserts (PICM0RG50; Millipore, Darmstadt, Germany), placed on the culture medium (50% minimal essential medium [MEM], 21% Hank’s balanced salt solution, 15 mM NaHCO_3_, 6.25 mM N-2-hydroxyethylpiperazine-N’-2-ethanesulfonic acid, 10 mM D-glucose, 1 mM L-glutamine, 0.88 mM ascorbic acid, 1 mg/mL insulin, and 25% horse serum), and incubated at 35° °C in 5% CO_2_.

For the imaging of spine morphology and sLTP, the cultured neurons were transfected by an injection of AAVs using a glass pipette (Narishige, Tokyo, Japan) after 2–6 days in the slice culture. For Ca^2+^ imaging, other cultured neurons were transfected with a gene gun (Scientz Biotechnology, Ningbo, China) using 1.6 μm gold particles coated with plasmids after 8–9 days in slice culture. To make the bullets for the gene gun, gold particles (6 mg) and DNA (12 μg) were used in a 30 cm long tube.

### The induction of homeostatic plasticity by pharmacological neuronal excitation

GABA_A_ receptor antagonists (10 μM bicuculline or 1–3 μM gabazine) were added to the culture medium after 11–13 days *in vitro* (DIV). The time of application was then set to 0 h for the experiments. The cultured hippocampal slices were incubated in this culture medium for 24 hrs at 35° C in 5% CO_2_. Subsequently, the slices were placed in an imaging buffer solution, and the experiment was carried out for up to 3 hrs.

### Imaging and analysis of spine morphology

Dendritic spine imaging of hippocampal slice cultures was performed using a custom two-photon microscope. A Ti: sapphire laser (Spectra-Physics, Santa Clara, CA, USA) tuned to 920 nm was used to excite the Achilles or d2Achilles proteins. The fluorescence signals of these proteins were collected with a ×60, NA1.0 objective lens (Olympus, Tokyo, Japan) and detected by a photomultiplier tube (H7422-40p; Hamamatsu, Hamamatsu, Japan) through an emission filter (FF01-510/84; Chroma). Signal acquisition and image (128×128 pixels) construction were carried out using a data acquisition board (PCI-6110; National Instruments, Austin, TX, USA) and ScanImage software (Pologruto et al., 2003).

To measure the spine density, three secondary dendrites (50 μm in length from the primary dendrite) were selected and analyzed for each neuron. The spine density was calculated by dividing the spine number by the length of the dendrite. The images were analyzed using ImageJ software (National Institute of Health, Bethesda, MD, USA).

### Two-photon glutamate uncaging

To induce sLTP in single spines, bath-applied 2 mM MNI-caged glutamate was uncaged by a second Ti: sapphire laser at a wavelength of 720 nm (30 trains, 0.5 Hz, 6 ms duration/pulse, 6 mW, measured under an objective lens) near the spine of interest. Since the focal plane of the imaging (920 nm) and activation (720 nm) lasers were different (~0.7 μm) due to chromatic aberration in the microscope, they were compensated by moving the sample stage along the z-axis (0.3 μm) with piezo stages (PKVL64F-100U; NCS6101C; Kohzu, Kawasaki, Japan) during the stimulation. Two-photon glutamate uncaging was carried out in the imaging buffer solution (136 mM NaCl, 5 mM KCl, 0.8 mM KH_2_PO_4_, 20 mM NaHCO_3_, 1.3 mM L-glutamine, 0.2 mM ascorbic acid, MEM amino acids solution [Gibco; Thermo Fisher, Waltham, MA, USA], MEM vitamin solution [Gibco; Thermo Fisher, Waltham, MA, USA], and 1.5 mg/ml phenol red) containing 4 mM CaCl_2_, 0 mM MgCl_2_, 1 μM tetrodotoxin, and 2 mM MNI-caged glutamate aerated with 95% O_2_/5% CO_2_ at 24–26°C.

### Ca^2+^ imaging in single spines

To measure the Ca^2+^ transients, a Ti: sapphire laser tuned to a wavelength of 1000 nm was used for the excitation of both GCaMP6f and mScarlet. For image acquisition, 128×32 pixels were acquired at 15.6 Hz. To apply the glutamate stimulation at single spines, bath-applied 2 mM MNI-caged glutamate was uncaged by a second Ti: sapphire laser at a wavelength of 720 nm (1 train, 6 ms duration/pulse, 6 mW, measured under an objective lens) near the spine of interest. Since the focal plane of the imaging (1000 nm) and activation (720 nm) lasers were different (0.5–1.0 μm), it was compensated by moving the sample stage along the z-axis (0.8 μm) with piezo stages during the stimulation. Two-photon glutamate uncaging was carried out in the imaging buffer solution containing 4 mM CaCl_2_, 0 mM MgCl_2_, 1 μM tetrodotoxin, and 2 mM MNI-caged glutamate aerated with 95% O_2_/5% CO_2_ at 24–26°C.

### Two-photon paCaMKII activation

To activate paCaMKII in single spines with two-photon excitation, a second Ti: sapphire laser tuned to a wavelength of 820 nm was used with 30 trains (0.5 Hz, 80 ms duration/pulse, 4 mW) in the spine of interest. Since the focal plane of the imaging (1010 nm) and activation (820 nm) lasers were different (0.5–1.0 μm), it was compensated by moving the sample stage in the z-axis (0.75 μm) with piezo stages during the stimulation. Two-photon paCaMKII activation was carried out in the imaging buffer solution containing 2 mM CaCl_2_ and 2 mM MgCl_2_ aerated with 95% O_2_/5% CO_2_ at 24–26°C.

### Primary neuronal culture and AAV infection

Low-density cultures of dissociated embryonic mouse hippocampal neurons were prepared as described previously (Murakoshi et al., 2017). Briefly, hippocampi were removed from C57BL/6N mice at embryonic day 18 and treated with papain for 10 min at 37°C, followed by gentle trituration. Hippocampal neurons were seeded onto polyethylenimine-coated 3-cm dishes (2 × 10^5^ cells/dish) and cultured in neurobasal medium (Gibco; Thermo Fisher, Waltham, MA, USA) supplemented with B-27 and 2 mM Glutamax (Gibco; Thermo Fisher, Waltham, MA, USA). Primary neuronal cultures were infected with AAV-ESARE-mScarlet-Flag particles at a titer of 4.35×10^6^ genome copies/mL and AAV-CaMK0.4-FHS-paCaMKII-WPRE3 at a titer of 4.72×10^6^ genome copies/ml at DIV 9. After ~65 hrs, we applied 10 μM bicuculline to the cultured neurons for 24 hrs, followed by a biochemical assay.

### Biochemical assay of autophosphorylation

For the paCaMKII autophosphorylation assay in the cultured hippocampal neurons, the neurobasal medium was replaced with the medium containing no bicuculline and incubated for 1 hour in a CO_2_ incubator in accordance with the condition of the sLTP experiment. To induce paCaMKII autophosphorylation, the samples were continuously illuminated with a light-emitting diode (M455L2-C1; Thorlabs, Newton, NJ, USA) at 3.82 mW cm^-2^ for 5 min. The reactions were stopped at the indicated times by adding a lysis solution (50 mM Tris pH 7.5, 1% NP-40, 5% glycerol, 150 mM NaCl, and 4 mM ethylenediaminetetraacetic acid). The samples were collected and centrifuged, and the supernatant was dissolved in sodium dodecyl sulfate sample buffer and analyzed by western blotting.

Western blotting was performed with the following antibodies: anti-phospho-CaMKII (Thr286) (D21E4; Cell Signaling Technology, MA, USA), anti-CaMKIIα (6G9; Cell Signaling Technology, MA, USA), anti-β-Actin (8H10D10; Cell Signaling Technology, MA, USA), anti-RFP for mScarlet detection (M204-3, MBL; Nagoya, Japan), and horseradish peroxidase-anti-mouse and -rabbit antibodies (Jackson Laboratory, Bar Harbor, ME, USA).

### Quantification and statistical analysis

Statistical analyses were performed using MATLAB (Math Works, MA, USA) or GraphPad Prism (GraphPad, SanDiego, CA, USA) software. The types of statistical tests, number of samples, and thresholds for statistical significance are described in the legends.

### Resource availability

Further information and requests for resources and reagents should be directed to and will be fulfilled by H.M.

## Notes

### Competing Interest Statement

The authors have declared no competing interest.

## REFERENCES

Abegg, M.H., Savic, N., Ehrengruber, M.U., McKinney, R.A., and Gahwiler, B.H. (2004). Epileptiform activity in rat hippocampus strengthens excitatory synapses. J Physiol 554, 439–448.

Bayer, K.U., and Schulman, H. (2019). CaM Kinase: Still Inspiring at 40. Neuron 103, 380–394.

Bindels, D.S., Haarbosch, L., van Weeren, L., Postma, M., Wiese, K.E., Mastop, M., Aumonier, S., Gotthard, G., Royant, A., Hink, M.A., et al. (2017). mScarlet: a bright monomeric red fluorescent protein for cellular imaging. Nat Methods 14, 53–56.

Bosch, M., Castro, J., Saneyoshi, T., Matsuno, H., Sur, M., and Hayashi, Y. (2014). Structural and molecular remodeling of dendritic spine substructures during long-term potentiation. Neuron 82, 444–459.

Chen, T.W., Wardill, T.J., Sun, Y., Pulver, S.R., Renninger, S.L., Baohan, A., Schreiter, E.R., Kerr, R.A., Orger, M.B., Jayaraman, V., et al. (2013). Ultrasensitive fluorescent proteins for imaging neuronal activity. Nature 499, 295–300.

Chen, Y., Wang, Y., Erturk, A., Kallop, D., Jiang, Z., Weimer, R.M., Kaminker, J., and Sheng, M. (2014). Activity-induced Nr4a1 regulates spine density and distribution pattern of excitatory synapses in pyramidal neurons. Neuron 83, 431–443.

Cingolani, L.A., and Goda, Y. (2008). Actin in action: the interplay between the actin cytoskeleton and synaptic efficacy. Nat Rev Neurosci 9, 344–356.

Cooper, L.N., and Bear, M.F. (2012). The BCM theory of synapse modification at 30: interaction of theory with experiment. Nat Rev Neurosci 13, 798–810.

Derkach, V.A., Oh, M.C., Guire, E.S., and Soderling, T.R. (2007). Regulatory mechanisms of AMPA receptors in synaptic plasticity. Nat Rev Neurosci 8, 101–113.

Donnelly, M.L., Luke, G., Mehrotra, A., Li, X., Hughes, L.E., Gani, D., and Ryan, M.D. (2001). Analysis of the aphthovirus 2A/2B polyprotein ‘cleavage’ mechanism indicates not a proteolytic reaction, but a novel translational effect: a putative ribosomal ‘skip’. J Gen Virol 82, 1013–1025.

Dorrbaum, A.R., Alvarez-Castelao, B., Nassim-Assir, B., Langer, J.D., and Schuman, E.M. (2020). Proteome dynamics during homeostatic scaling in cultured neurons. Elife 9.

Ehlers, M.D. (2003). Activity level controls postsynaptic composition and signaling via the ubiquitin-proteasome system. Nat Neurosci 6, 231–242.

Fiore, R., Rajman, M., Schwale, C., Bicker, S., Antoniou, A., Bruehl, C., Draguhn, A., and Schratt, G. (2014). MiR-134-dependent regulation of Pumilio-2 is necessary for homeostatic synaptic depression. EMBO J 33, 2231–2246.

Giese, K.P., and Mizuno, K. (2013). The roles of protein kinases in learning and memory. Learn Mem 20, 540–552.

Goold, C.P., and Nicoll, R.A. (2010). Single-cell optogenetic excitation drives homeostatic synaptic depression. Neuron 68, 512–528.

Govindarajan, A., Israely, I., Huang, S.Y., and Tonegawa, S. (2011). The dendritic branch is the preferred integrative unit for protein synthesis-dependent LTP. Neuron 69, 132–146.

Harvey, C.D., and Svoboda, K. (2007). Locally dynamic synaptic learning rules in pyramidal neuron dendrites. Nature 450, 1195–1200.

Herring, B.E., and Nicoll, R.A. (2016). Long-Term Potentiation: From CaMKII to AMPA Receptor Trafficking. Annu Rev Physiol 78, 351–365.

Hu, J.H., Park, J.M., Park, S., Xiao, B., Dehoff, M.H., Kim, S., Hayashi, T., Schwarz, M.K., Huganir, R.L., Seeburg, P.H., et al. (2010). Homeostatic scaling requires group I mGluR activation mediated by Homer1a. Neuron 68, 1128–1142.

Jourdain, P., Fukunaga, K., and Muller, D. (2003). Calcium/calmodulin-dependent protein kinase II contributes to activity-dependent filopodia growth and spine formation. J Neurosci 23, 10645–10649.

Kawashima, T., Kitamura, K., Suzuki, K., Nonaka, M., Kamijo, S., Takemoto-Kimura, S., Kano, M., Okuno, H., Ohki, K., and Bito, H. (2013). Functional labeling of neurons and their projections using the synthetic activity-dependent promoter E-SARE. Nat Methods 10, 889–895.

Keck, T., Hubener, M., and Bonhoeffer, T. (2017). Interactions between synaptic homeostatic mechanisms: an attempt to reconcile BCM theory, synaptic scaling, and changing excitation/inhibition balance. Curr Opin Neurobiol 43, 87–93.

Lee, K.J., Lee, Y., Rozeboom, A., Lee, J.Y., Udagawa, N., Hoe, H.S., and Pak, D.T. (2011). Requirement for Plk2 in orchestrated ras and rap signaling, homeostatic structural plasticity, and memory. Neuron 69, 957–973.

Lee, M.C., Yasuda, R., and Ehlers, M.D. (2010). Metaplasticity at single glutamatergic synapses. Neuron 66, 859–870.

Lee, S.J., Escobedo-Lozoya, Y., Szatmari, E.M., and Yasuda, R. (2009). Activation of CaMKII in single dendritic spines during long-term potentiation. Nature 458, 299–304.

Li, X., Zhao, X., Fang, Y., Jiang, X., Duong, T., Fan, C., Huang, C.C., and Kain, S.R. (1998). Generation of destabilized green fluorescent protein as a transcription reporter. J Biol Chem 273, 34970–34975.

Lisman, J., Yasuda, R., and Raghavachari, S. (2012). Mechanisms of CaMKII action in long-term potentiation. Nat Rev Neurosci 13, 169–182.

Lledo, P.M., Hjelmstad, G.O., Mukherji, S., Soderling, T.R., Malenka, R.C., and Nicoll, R.A. (1995). Calcium/calmodulin-dependent kinase II and long-term potentiation enhance synaptic transmission by the same mechanism. Proc Natl Acad Sci U S A 92, 11175–11179.

Lock, M., Alvira, M., Vandenberghe, L.H., Samanta, A., Toelen, J., Debyser, Z., and Wilson, J.M. (2010). Rapid, simple, and versatile manufacturing of recombinant adeno-associated viral vectors at scale. Hum Gene Ther 21, 1259–1271.

Malinow, R., and Malenka, R.C. (2002). AMPA receptor trafficking and synaptic plasticity. Annu Rev Neurosci 25, 103–126.

Matsuzaki, M., Honkura, N., Ellis-Davies, G.C., and Kasai, H. (2004). Structural basis of long-term potentiation in single dendritic spines. Nature 429, 761–766.

Mendez, P., Stefanelli, T., Flores, C.E., Muller, D., and Luscher, C. (2018). Homeostatic Plasticity in the Hippocampus Facilitates Memory Extinction. Cell Rep 22, 1451–1461.

Moulin, T.C., Petiz, L.L., Rayee, D., Winne, J., Maia, R.G., Lima da Cruz, R.V., Amaral, O.B., and Leao, R.N. (2019). Chronic in vivo optogenetic stimulation modulates neuronal excitability, spine morphology, and Hebbian plasticity in the mouse hippocampus. Hippocampus 29, 755–761.

Murakoshi, H., Shin, M.E., Parra-Bueno, P., Szatmari, E.M., Shibata, A.C., and Yasuda, R. (2017). Kinetics of Endogenous CaMKII Required for Synaptic Plasticity Revealed by Optogenetic Kinase Inhibitor. Neuron 94, 37–47 e35.

Murakoshi, H., and Yasuda, R. (2012). Postsynaptic signaling during plasticity of dendritic spines. Trends Neurosci 35, 135–143.

Nakahata, Y., and Yasuda, R. (2018). Plasticity of Spine Structure: Local Signaling, Translation and Cytoskeletal Reorganization. Front Synaptic Neurosci 10, 29.

Nicoll, R.A. (2017). A Brief History of Long-Term Potentiation. Neuron 93, 281–290.

Perez-Otano, I., and Ehlers, M.D. (2005). Homeostatic plasticity and NMDA receptor trafficking. Trends Neurosci 28, 229–238.

Pettit, D.L., Perlman, S., and Malinow, R. (1994). Potentiated transmission and prevention of further LTP by increased CaMKII activity in postsynaptic hippocampal slice neurons. Science 266, 1881–1885.

Pologruto, T.A., Sabatini, B.L., and Svoboda, K. (2003). ScanImage: flexible software for operating laser scanning microscopes. Biomed Eng Online 2, 13.

Sala, C., Futai, K., Yamamoto, K., Worley, P.F., Hayashi, Y., and Sheng, M. (2003). Inhibition of dendritic spine morphogenesis and synaptic transmission by activity-inducible protein Homer1a. J Neurosci 23, 6327–6337.

Saneyoshi, T., Matsuno, H., Suzuki, A., Murakoshi, H., Hedrick, N.G., Agnello, E., O’Connell, R., Stratton, M.M., Yasuda, R., and Hayashi, Y. (2019). Reciprocal Activation within a Kinase-Effector Complex Underlying Persistence of Structural LTP. Neuron 102, 1199–1210 e1196.

Schanzenbacher, C.T., Langer, J.D., and Schuman, E.M. (2018). Time- and polarity-dependent proteomic changes associated with homeostatic scaling at central synapses. Elife 7.

Schanzenbacher, C.T., Sambandan, S., Langer, J.D., and Schuman, E.M. (2016). Nascent Proteome Remodeling following Homeostatic Scaling at Hippocampal Synapses. Neuron 92, 358–371.

Seeburg, D.P., and Sheng, M. (2008). Activity-Induced Polo-Like Kinase 2 Is Required for Homeostatic Plasticity of Hippocampal Neurons during Epileptiform Activity. The Journal of Neuroscience 28, 6583–6591.

Shibata, A.C.E., Ueda, H.H., Eto, K., Onda, M., Sato, A., Ohba, T., Nabekura, J., and Murakoshi, H. (2021). Photoactivatable CaMKII induces synaptic plasticity in single synapses. Nat Commun 12, 751.

Sobczyk, A., Scheuss, V., and Svoboda, K. (2005). NMDA receptor subunit-dependent [Ca2+] signaling in individual hippocampal dendritic spines. J Neurosci 25, 6037–6046.

Stoppini, L., Buchs, P.A., and Muller, D. (1991). A simple method for organotypic cultures of nervous tissue. J Neurosci Methods 37, 173–182.

Suarez, L.M., Cid, E., Gal, B., Inostroza, M., Brotons-Mas, J.R., Gomez-Dominguez, D., de la Prida, L.M., and Solis, J.M. (2012). Systemic injection of kainic acid differently affects LTP magnitude depending on its epileptogenic efficiency. PLoS One 7, e48128.

van der Linden, J.A., Joels, M., Karst, H., Juta, A.J., and Wadman, W.J. (1993). Bicuculline increases the intracellular calcium response of CA1 hippocampal neurons to synaptic stimulation. Neurosci Lett 155, 230–233.

Watt, A.J., van Rossum, M.C., MacLeod, K.M., Nelson, S.B., and Turrigiano, G.G. (2000). Activity coregulates quantal AMPA and NMDA currents at neocortical synapses. Neuron 26, 659–670.

Yashiro, K., and Philpot, B.D. (2008). Regulation of NMDA receptor subunit expression and its implications for LTD, LTP, and metaplasticity. Neuropharmacology 55, 1081–1094.

Yoshioka-Kobayashi, K., Matsumiya, M., Niino, Y., Isomura, A., Kori, H., Miyawaki, A., and Kageyama, R. (2020). Coupling delay controls synchronized oscillation in the segmentation clock. Nature 580, 119–123.

